# Two IncHI1 megaplasmids in *Klebsiella* species reveal transposable-element-mediated *bla*IMP-1 mobilisation

**DOI:** 10.64898/2026.09.09.750360

**Authors:** Yu Wan, Victoria Orr, Maria Getino, Sophie Mannix, Chloe Heenan, Ebony Richmond-Mensah, Nicholas Harper, Joshua L. C. Wong, Rojus Urbonas, Martina O. Chukwu, Jane F. Turton, Katie L. Hopkins, Gad Frankel, Alison H. Holmes, Frances Davies, Elita Jauneikaite

## Abstract

**Introduction:** Carbapenemase-producing *Enterobacterales* (CPE) represent a major threat to hospitalised patients worldwide. The dissemination of carbapenemase genes, such as *bla*_IMP_, is frequently mediated by mobile genetic elements including plasmids. During a previously described multispecies, healthcare-associated outbreak of *bla*_IMP_-positive CPE in North West London, two unusual isolates, IMP47 (*Klebsiella grimontii*) and IMP76 (*K. pneumoniae*), recovered in 2019 from routinely collected rectal swabs of inpatients, were predicted to harbour *bla*_IMP-1_-carrying IncHI1 megaplasmids.

**Aims:** This study aimed to determine complete genomic sequences of IMP47 and IMP76, resolve the genetic context of *bla*_IMP-1_, and assess the conjugative mobility of *bla*_IMP-1_-carrying megaplasmids.

**Methods:** Genomic sequences of both isolates were recovered through hybrid assembly of Oxford Nanopore and Illumina sequencing reads. Complete plasmid sequences were characterised to determine replicons, conjugation machinery, and genes encoding resistance to antimicrobials or other stress factors. Integrons and transposable elements (TEs) within flanking regions of *bla*_IMP-1_ were resolved through genome annotation and search against public databases. Liquid-mating experiments were performed to assess the mobility of *bla*_IMP-1_-carrying plasmids.

**Results:** Completed genome assemblies were generated from both isolates, confirming two *bla*_IMP-1_-carrying megaplasmids, pIMP47 (391 kbp) and pIMP76_1 (519 kbp), of the replicon type IncHI1A(pNDM-CIT)/IncHI1B(pNDM-CIT). The *bla*_IMP-1_ locus was carried by nearly identical class 1 integrons in both plasmids and a closely related IncHI1 megaplasmid pEB3_IMP1 (361 kbp) previously identified in South West England. Comparative analysis revealed conserved genetic structures linking *bla*_IMP-1_ to mercury-resistance genes and TEs Tn*c025*, Tn*As3*, IS*26*, and IS*5075*, suggesting a history of recombination and potential for TE-mediated mobilisation. Conjugation experiments confirmed transfer of pIMP76_1 into a recipient *K. pneumoniae* strain, resulting in acquisition of ertapenem resistance, whereas transfer of pIMP47 was not observed under the tested conditions.

**Conclusion:** Two IMP-producing IncHI1 megaplasmids in gut-colonising *Klebsiella* species revealed TE-mediated *bla*_IMP-1_ mobilisation. The co-localisation of *bla*_IMP-1_ and metal-resistance genes in both plasmids highlights the potential for co-selection of *bla*_IMP-1_ in environments enriched with metal ions. Our findings underscore the importance of longitudinal genomic surveillance of carbapenemase-encoding megaplasmids in healthcare settings.

**Impact statement:** This study characterises two *bla*_IMP-1_-carrying IncHI1 megaplasmids in *Klebsiella* isolates recovered in 2019 from rectal swabs of inpatients in London. Comparative genomic analysis revealed that *bla*_IMP-1_ was embedded within a conserved class 1 integron linked to insertion sequences and transposon-borne mercury-resistance genes, suggesting that transposable-element-mediated recombination has contributed to the acquisition of *bla*_IMP-1_ in IncHI1 plasmids and may drive further mobilisation of this carbapenem-resistance determinant. Experimental evidence confirmed conjugative transfer of one megaplasmid and associated ertapenem resistance encoded by this plasmid. Our findings identify IncHI1 megaplasmids as potential emerging vectors of *bla*_IMP-1_ and emphasise the importance of genomic surveillance for monitoring the spread of carbapenemase-encoding megaplasmids in healthcare settings.

**Data summary:** Raw Illumina and nanopore whole-genome sequencing reads of isolates IMP47 and IMP76 have been deposited in the European Nucleotide Archive (ENA, www.ebi.ac.uk/ena) under BioSample accessions SAMEA6990777 and SAMEA6990795, respectively. Complete genome assemblies of these isolates are accessible in the ENA under Assembly accessions GCA_903936125.2 (IMP47) and GCA_903936275.2 (IMP76). All data accessions are available in supplementary Table S1.

## Introduction

Antimicrobial resistance (AMR) is a major global threat to human health [1], and carbapenems are among the last-resort antimicrobials used for treating multidrug-resistant bacterial infections [2]. In particular, infections caused by carbapenemase-producing *Enterobacterales* (CPE) present substantial clinical and economic challenges [3]. AMR genes, including those encoding carbapenemases such as imipenemase (IMP), are frequently localised on mobile genetic elements (MGEs), which facilitate their dissemination both within and between bacterial species [4, 5].

Discovered in a *P. aeruginosa* clinical isolate collected in Japan in 1988 [6], IMP is a family of zinc-dependent metallo-β-lactamases in Ambler class B [7]. IMP carbapenemases can effectively hydrolyse all available β-lactam antimicrobials (except aztreonam) and are refractory to clinically available β-lactamase inhibitors such as tazobactam, clavulanic acid, avibactam, relebactam, and vaborbactam [8–11]. A study of 4,556 publicly available, *bla*_IMP_-containing bacterial genomes isolated between 1996 and 2023 across the globe revealed IncHI2 plasmids as a predominant vector [12]. Consisting of two major groups, IncHI1 and IncHI2, the majority of IncH plasmids can be referred to as megaplasmids— exceptionally large plasmids of lengths greater than 100–350 kbp [13, 14]. Megaplasmids have been widely recognised as drivers of multidrug resistance (MDR) due to their flexible capacity to co-mobilise diverse AMR genes [15–17]. Moreover, these plasmids frequently carry metal-resistance genes that facilitate co-selection of AMR and drive niche adaptation [16, 18]. This dual resistance profile exacerbates the persistence of bacterial pathogens in clinical and environmental reservoirs [14, 16].

IMP-encoding IncH megaplasmids, mostly IncHI2, were identified in isolates collected from patients in London, UK, between June 2016 and November 2019 during an investigation of a regional, multispecies outbreak of CPE carrying both *bla*_IMP_ and *mcr-9* genes [19]. Two of these isolates, *Klebsiella grimontii* IMP47 in the *K. oxytoca* species complex (*Ko*SC) and *K. pneumoniae* IMP76, carried a *bla*_IMP_ variant *bla*_IMP-1_, which was predicted to be localised on two putative IncHI1 megaplasmids according to draft genome assemblies [19]. A literature search in the PubMed (pubmed.ncbi.nlm.nih.gov) [query: ((“IMP”) OR (“blaIMP”)) AND (“IncHI1”)] and Scopus (www.scopus.com) [query: (ALL(“blaIMP”) OR ALL(“IMP”)) AND ALL(“IncHI1”)] databases on 5 August 2026 indicated no previous report on this *bla*_IMP-1_-IncHI1 association. In this study, we aim to recover and characterise complete sequences of these *bla*_IMP-1_-carrying IncHI1 plasmids by leveraging hybrid genome assembly.

## Methods

### Isolate collection

IMP47 and IMP76 of *Klebsiella* species were recovered from routinely collected rectal swabs of two inpatients in April and January 2019, respectively, in two hospitals of Imperial College Healthcare National Health Service (NHS) Trust in North West London. As per a local policy for enhanced CPE screening [20], both patients were identified for their recent or current stays in these sub-regions of London, and neither patient was screened previously. Rectal swabs of these patients were sent to North West London Pathology for microbiological characterisation, and the isolates were collected as part of an investigation of healthcare-associated IMP-producing CPE [19].

### Whole-genome sequencing

IMP47 and IMP76 were aerobically cultivated on Luria-Bertani agar (Merck, Germany) at 37°C overnight. Genomic DNA was extracted from each culture using GenElute Bacterial Genomic DNA Kit NA2110 (Merck) following the manufacturer’s instructions. The nanopore sequencing was performed using the Rapid Barcoding Kit SQK-RBK114.24 and MinION R10.4.1 flow cell FLO-MIN114 (Oxford Nanopore Technologies, UK). Super-accuracy basecalling of the nanopore data was conducted on the Dorado basecalling server v7.8.3 (model: dna_r10.4.1_e8.2_400bps_sup@v4.3.0) through MinKNOW v25.05.14 (Oxford Nanopore Technologies). Previously generated Illumina sequencing data of these two isolates were downloaded from the National Center for Biotechnology Information (NCBI) Sequence Read Archive (accessions: IMP47, ERR4280191; IMP76, ERR4280209) [19] using fastq-dump in NCBI’s SRA-Toolkit v3.2.1 (github.com/ncbi/sra-tools).

### Quality control of sequencing reads

Read quality was assessed using FastQC v0.12.1 [21], followed by platform-specific steps (Table S2): Illumina reads were additionally assessed with MultiQC v1.28 [22] before trimming and filtering with fastp v0.26.0 [23], while nanopore reads were additionally assessed with NanoPlot v1.44.1 [24] before trimming and filtering with fastplong v0.3.0 [23]. The processed nanopore reads were further filtered for a minimum average quality of Phred Q15 for IMP47 and Q10 for IMP76 using nanoq v0.10.0 [25].

To evaluate the level of contamination, taxonomical profiles of processed Illumina and nanopore reads were assessed using Kraken2 v2.14 [26] and a pre-built full Standard database (benlangmead.github.io/aws-indexes/k2; released on 2 April 2025), and the taxonomical abundance was estimated from Illumina reads and Kraken2 outputs using Bracken v3.1 [27].

### Hybrid genome assembly

To decide long-read assemblers, draft assemblies of IMP47 and IMP76 genomes were generated from processed nanopore reads using Flye v2.9.6 [28], and the read depth of each assembly was estimated by dividing the total read length (determined using Seqkit v2.10.0 [29]) with the assembly length as per the Lander-Waterman equation [30]. Formal genome assemblies of IMP47 and IMP76 were generated from processed nanopore reads using Trycycler v0.5.5 [31] and Flye v2.9.6, respectively, according to estimated read depths (IMP47: 76×; IMP76: 34×) and correspondingly adjusted parameters. These assemblies were polished with nanopore reads using Medaka v2.1.0 (Oxford Nanopore Technologies) followed by three rounds of Illumina-read polishing, for which the first and final rounds were completed with Polypolish v0.6.0 [32] and the second round with Pypolca v0.3.1 [33]. Moreover, circularised contigs in the assembly of IMP76 underwent reorientation using dnaapler v1.2.0 [34] between the second and third round of polishing, whereas circularised contigs in the assembly of IMP47 were reoriented by Trycycler [31]. Depths of processed Illumina and nanopore reads were confirmed using the final assemblies and mosdepth v0.3.11 [35].

### Genetic characterisation

Complete assemblies of IMP47 and IMP76 genomes were annotated using bakta v1.11.3 and its full database v6.0 (zenodo.org/records/14916843) [36]. Plasmid replicon types were detected using Abricate v1.2.0 and the PlasmidFinder database (updated on 19 December 2025) for a minimum nucleotide identity and reference coverage of 80% [37, 38]. A perfect match between a plasmid of interest and a known replicon type was identified when corresponding sequence probes in the PlasmidFinder database completely aligned (100% nucleotide identity and coverage) to the plasmid. Furthermore, plasmid mobility, relaxase types, and mating pair formation (MPF) were predicted using mob_typer of MOBsuite v3.1.9 [39]; the origin of replication (*ori*) regions were identified using bakta, and the origin of transfer (*oriT*) sites were identified using OriTFinder2 (bioinfo-mml.sjtu.edu.cn/oriTDB2/oriTfinder.php, accessed on 18 October 2025) [40]. Genes coding for conjugation machinery were identified using both bakta annotations and mob_typer’s outputs.

Genes coding for AMR and stress responses (heavy metal, biocide, and heat) in IMP47 and IMP76 were identified using AMRFinderPlus v4.0.23 and its database v2025-07-16.1 (Table S2) [41] and were checked with genome annotations. Transposable elements (TEs) were searched against the ISfinder and TnCentral databases using nucleotide BLAST implemented on each database’s website [42, 43]. Integrons were detected using IntegronFinder v2.0.5 installed on the European Galaxy Server [44, 45]. Multi-locus sequence typing (MLST) for IMP47 was completed using PubMLST’s online sequence query tool (pubmlst.org) and the *Ko*SC database. The MLST for IMP76 was completed using the same tool, hosted by the Pasteur Institute (bigsdb.pasteur.fr), and the *K. pneumoniae* species complex (*Kp*SC) database. Capsule and O-antigen types were predicted from genome assemblies using Kleborate v3.2.4 [46, 47].

### Comparative analysis

Plasmid sequences from IMP47 and IMP76 were searched in the *Klebsiella* (taxid: 570), *Enterobacter* (taxid: 547), *Escherichia* (taxid: 561), *Shigella* (taxid: 620), *Salmonella* (taxid: 590), *Citrobacter* (taxid: 544), and plasmids (taxid: 36549) collections of NCBI’s core nucleotide database (core_nt) using megaBLAST (blast.ncbi.nlm.nih.gov, accessed on 27 August 2026) to identify homologues. The closest matching *bla*_IMP_-carrying IncHI1 plasmids, along with the 289-kbp reference IncHI1 plasmid pNDM-CIT (GenBank accession: JX182975.1) in the PlasmidFinder database, were downloaded from GenBank for comparison. All downloaded plasmid sequences were reorientated with dnaapler, reannotated with bakta, and genetically characterised in the same ways as IMP47 and IMP76 to avoid technical discrepancies. Comparison between IncHI1 plasmids were performed using BLASTn (E-value ≤0.0001, with low-complexity regions masked) as implemented on the ProkSee webserver (proksee.ca) [48], cross-checked with results from the command-line megaBLAST (nucleotide identity ≥80%; E-value ≤0.0001). Both megaBLAST and BLASTn were components of BLAST+ v2.16.0 [49]. Genomic comparison and gene synteny were visualised using Proksee and the R package gggenomes, respectively [48, 50].

To investigate the origin of a putatively exogenous IncR region in megaplasmid pIMP76_1, the nucleotide sequence of this region was searched against the same sequence collections of the NCBI core_nt database as those for the plasmids from IMP47 and IMP76 using megaBLAST (accessed on 27 August 2026). Resulting alignments were filtered for nucleotide identities greater than 99% and ranked by query coverages to identify the closest match, for which plasmid replicon types were detected using the PlasmidFinder webserver v3.0.3 (genepi.dk/plasmidfinder) with its *Enterobacterales* database v2.2.0 (minimum nucleotide identity: 90%; minimum coverage: 80%). Moreover, this IncR region was searched against the plasmid database PLSDB v2024_05_31_v2 [51] using BLASTn in BLAST+ v2.16.0 with parameters ‘-perc_identity 50 -qcov_hsp_perc 20’ [49].

Gene content and genetic structure of *bla*_IMP-1_-carrying plasmids were compared using a pangenome approach implemented in Panaroo v1.7.0 [52], which took as input bakta annotations of plasmids and produced a pangenome graph and gene presence-absence profiles under the sensitive mode and default identity thresholds (Table S2). The pangenome graph was visualised as a network in the yFiles Organic layout using Cytoscape Desktop v3.10.4 [53]. Type-F mating-pair stabilisation protein TraN and its IncHI counterpart TrhN, determinants of plasmid species specificity, were compared across *bla*_IMP-1_-carrying plasmids within the context of previously reported long, medium, short, and V-shaped types and α–δ subtypes of TraN (Table 1) [54, 55]. For comparison within each collection of TraN and TrhN sequences (for instance, type-specific or multi-type), a stratified ensemble of 4×4 (‘-replicates 4’) multi-sequence global alignments were generated using MUSCLE v5.3 [56], and the ensemble was considered robust if its letter-pair dispersion (D_LP) was below 0.05. For each robust ensemble, a maximum phylogenetic tree was reconstructed from the most confident global alignment with IQ-Tree v3.1.3 (Table S2) and visualised with iTOL v7 [57, 58]. Three-dimensional structures of TraN/TrhN were predicted using AlphaFold3 [59] as implemented on AlphaFolder Server (alphafoldserver.com; accessed on 30 July 2026), and the most confident model (Rank 0) of each protein was visualised in ChimeraX v1.12 [60].

**Table 1.** TraN sequences used for comparison. The long (L), medium (M), short (S), and V-shaped (V) types and subtypes (α–δ) of TraN were previously defined [54, 55]. Accessions refer to sequence data in the NCBI Protein database, and the length of each protein is measured by the number of amino acids (aa). Accessions of TrhN_1_, TrhN_2_, and TraN_S1_ were determined by searching for identical protein sequences in the NCBI RefSeq Protein database for *Enterobacterales* (taxid: 91347) with BLASTp. Plasmids and hosts refer to sources where the TraN proteins were reported. Plasmids indicated by asterisks are reported in this study. TrhN_1_ was identical to the TrhN (accession: WRP35066.1) produced by pIMP3_IMP1, originally reported in *Enterobacter hormaechei*.

| TraN | Type | Subtype | Accession | Length (aa) | Plasmid | Host |
| --- | --- | --- | --- | --- | --- | --- |
| TraN<sub>S</sub>α1 | S | α | ABD60034.1 | 616 | R100-1 | *Escherichia coli* |
| TraN<sub>S</sub>α2 | S | α | AAL23498.1 | 608 | pSLT | *Salmonella enterica* |
| TraN<sub>S</sub>β1 | S | β | ARQ19727.1 | 651 | pKpQIL-UK | *Klebsiella pneumoniae* |
| TraN<sub>S</sub>β2 | S | β | BAS44060.1 | 651 | pKO_JKo3_3 | *Klebsiella oxytoca* |
| TraN<sub>S</sub>γ | S | γ | WP_000821835.1 | 602 | F | *Enterobacteriaceae* spp. |
| TraN<sub>S</sub>δ1 | S | δ | ANZ89826.1 | 617 | pOZ172 | *Citrobacter freundii* |
| TraN<sub>S</sub>δ2 | S | δ | WP_001398575.1 | 612 | pCss165Kan | *Escherichia coli* |
| TraN<sub>M</sub>α | M | α | ACN66968.1 | 912 | pRA1 | *Aeromonas hydrophila* |
| TraN<sub>M</sub>β | M | β | AHI38890.1 | 931 | pNDM-US | *Klebsiella pneumoniae* |
| TraN<sub>L</sub>α | L | α | AAF69844.1 | 1058 | R27 | *Salmonella enterica* |
| TraN<sub>L</sub>β | L | β | APZ78090.1 | 1062 | pHNAH67 | *Escherichia coli* |
| TraN<sub>L</sub>γ | L | γ | AFB82831.1 | 1061 | pNDM-MAR | *Klebsiella pneumoniae* |
| TraN<sub>V</sub>α | V | α | QBN23304.1 | 891 | pABAY10001_1C | *Acinetobacter baumannii* |
| TrhN<sub>1</sub> | L | Unknown | WRP35066.1 | 1058 | pIMP47 * | *Klebsiella grimontii* |
| TrhN<sub>2</sub> | L | Unknown | WP_251894854.1 | 1058 | pIMP76_1 * | *Klebsiella pneumoniae* |
| TraN<sub>S1</sub> | S | Unknown | WP_023287134.1 | 651 | pIMP76_1 * | *Klebsiella pneumoniae* |

### Antimicrobial susceptibility testing

Ertapenem and rifampicin minimum inhibitory concentrations (MICs) of IMP47, IMP76, and *K. pneumoniae* laboratory strain ICC8001 — a plasmid recipient with a rifampicin MIC above 100 mg/L [61] — were determined in technical triplicates using broth microdilution (BMD) as per the European Committee on Antimicrobial Susceptibility Testing (EUCAST) guidelines [40]. *Escherichia coli* strain ATCC 25922 and *Staphylococcus aureus* ATCC 29213 were used as quality controls for the ertapenem and rifampicin BMD, respectively [62]. Ertapenem MICs were interpreted with the EUCAST clinical breakpoint v16, whereas no rifampicin clinical breakpoint was available for *Enterobacterales* [41].

### Conjugation experiment

IMP47, IMP76, and ICC8001 were grown on cation-adjusted Muller-Hinton (MH2) agar (MH2 broth: 90922-500G, Millipore; microbiological agar: LP0011B, Oxoid) at 37 °C overnight, and then a single colony of each isolate was transferred into 10 mL MH2 broth and was incubated at 37 °C with shaking at 200 revolutions per minute (RPM) for three hours to reach the mid-log phase. Optical density (OD) was measured for each broth culture at the 600-nm wavelength in triplicates of 100 μL with a microplate (655161, Greiner Bio-One, UK) and a Multiskan FC 96-well plate photometer (ThermoFisher Scientific). ODs of paired donor and recipient cultures were equalised with MH2 broth. Then, liquid mating was performed by mixing 2.15 mL culture of a donor (IMP47 or IMP76) with 2.15 mL culture of the recipient (ICC8001) in a 50-mL centrifuge tube. The start-point OD of each mixture was immediately measured in triplicates of 100 μL, leaving 4 mL mixture for incubation. Control cultures were created by transferring 4 mL of IMP47, IMP76, and ICC8001, respectively, into 50-mL centrifuge tubes. The resulting five cultures (two mating mixture and three controls) were aerobically incubated at 26 °C [63] for six hours with gentle shaking at 60 RPM.

At the end of the mating period, the end-point OD was measured in 100-μL triplicates for each culture, and each mating mixture was spread onto five MH2 agar plates (100 μL per plate) containing 64 mg/L rifampicin and 0.5 mg/L ertapenem (rifampicin–ertapenem plates) to select for transconjugants. Moreover, 100 μL of each culture (mating mixture or control) was spread on an MH2 agar plate containing either 0.5 mg/L ertapenem (ertapenem-only plate) or 64 mg/L rifampicin (rifampicin-only plate) or without any antimicrobial for quality control. Inoculated plates were incubated at 37 °C overnight. Transconjugant colonies from each mating mixture were then counted, and three such colonies were subjected to the same rifampicin and ertapenem BMD as that for the donors and recipient. String tests were conducted to further distinguish between the recipient, donors, and selected transconjugants when cultured on MH2 agar. A positive test result or hypermucoviscous isolate was defined as the formation of a viscous string exceeding 5 mm in length [42].

## Results

### Ertapenem and rifampicin susceptibility of isolates

IMP47 and IMP76 were ertapenem-resistant, with MICs of 2 mg/L and >2 mg/L, respectively, while ICC8001 was ertapenem-susceptible (MIC: 0.03 mg/L). The rifampicin MIC of ICC8001 was >512 mg/L, while those of IMP47 and IMP76 were 16 mg/L and 32 mg/L, respectively.

### Genomic characteristics

IMP47 and IMP76 belonged to *Ko*SC ST564 and *Kp*SC ST1310, respectively. IMP47 had a chromosome size of 5,752,224 bp and carried a 391,260-bp, *bla*_IMP-1_-carrying megaplasmid pIMP47 (Figure S1). IMP76 had a chromosome size of 5,245,774 bp and carried two megaplasmids, pIMP76_1 (*bla*_IMP-1_-carrying, Figure S2) and pIMP76_2, with sizes of 518,625 bp and 208,755 bp, respectively. IMP47 and IMP76 harboured stress response genes (Table 2), with each isolate’s chromosome coding for cation-efflux transporter FieF, which is associated with resistance to iron, zinc, manganese, and cadmium [64, 65].

**Table 2.** Summary of genomic characteristics of isolates IMP47 and IMP76 with GenBank accessions noted beneath molecular names. Determinants of antimicrobial resistance (AMR) and stress responses were identified using both AMRFinderPlus and bakta. The phenotype associated with each determinant is indicated by brackets.

| Isolate | DNA molecule | Length (bp) | Replicon haplotype | Predicted mobility | AMR determinants | Stress response determinants |
| --- | --- | --- | --- | --- | --- | --- |
| IMP47<br>(*K. grimontii*) | Chromosome (OZ478279) | 5,752,224 | Not applicable | Not applicable | *emrD* (multidrug resistance)<br>*kdeA* (multidrug resistance)<br>*bla*<sub>OXY-6-1</sub> (extended-spectrum β-lactam resistance)<br>*fosA7* (fosfomycin resistance)<br>*oqxAB* (phenicol and quinolone resistance) | *fieF* (multi-metal resistance)<br>*ars*(*B, C, R*) (arsenic resistance) |
|  | pIMP47 (OZ478280) | 391,260 | IncHI1A/IncHI1B | Conjugative | *aac*(3)-*Ile* (aminoglycoside resistance)<br>*aac*(6')-*Ib4* (aminoglycoside resistance)<br>*bla*<sub>DHA-1</sub> (extended-spectrum β-lactam resistance)<br>*bla*<sub>IMP-1</sub> (carbapenem resistance)<br>*mph*(*A*) (macrolide resistance)<br>*qnrB4* (quinolone resistance)<br>*sul1* (sulphonamide resistance)<br>*dfrA17* (trimethoprim resistance) | *ter*(*A, B, C, D, E, F, W, X, Y, Z*) (tellurium resistance)<br>*mer*(*A, C, D, E, P, R, T*) (mercury resistance)<br>*chrA* (chromium resistance)<br>*qacE* (quaternary ammonium resistance)<br>*qacEdelta1* (quaternary ammonium resistance) |
| IMP76<br>(*K. pneumoniae*) | Chromosome (OZ478276) | 5,245,774 | Not applicable | Not applicable | *emrD* (multidrug resistance)<br>*kdeA* (multidrug resistance)<br>*bla*<sub>SHV-1</sub> (broad-spectrum β-lactam resistance)<br>*fosA* (fosfomycin resistance)<br>*oqxAB* (phenicol and quinolone resistance) | *fieF* (multi-metal resistance) |
|  | pIMP76_1 (OZ478277) | 518,625 | IncHI1A/IncHI1B/IncR | Conjugative | *aac*(6')-*Ib4* (aminoglycoside resistance)<br>*bla*<sub>IMP-1</sub> (carbapenem resistance)<br>*sul1* (sulphonamide resistance) | *ter*(*A, B, C, D, E, F, W, X, Y, Z*) (tellurium resistance)<br>*mer*(*A, C, D, E, P, R, T*) (mercury resistance)<br>*qacEdelta1* (quaternary ammonium resistance) |
|  | pIMP76_2 (OZ478278) | 208,755 | IncFIB(K)/RepB/repFIB | Non-mobilisable | Not detected | *ars*(*A, B, C, D, H, R*) (arsenic resistance)<br>*pco*(*A, B, C, D, E, R, S*) (copper resistance)<br>*sil*(*A, B, C, E, F, P, R, S*) (copper and silver resistance)<br>*hsp20* (heat resistance)<br>*clpK1* (heat resistance)<br>*shsP* (heat resistance)<br>*yfdX*(1, 2) (heat resistance)<br>*hdeD-GI* (heat resistance)<br>*trxLHR* (heat resistance)<br>*kefB-GI* (heat resistance)<br>*psi-GI* (heat resistance) |

Both pIMP47 and pIMP76_1 were predicted to be conjugative and exhibited an average nucleotide identity of >99%, covering 94% and 71% of their lengths, respectively (Figure 1). The same *ori* region spanning positions 64,694–65,831 in pIMP47 and 209,696–210,833 in pIMP76_1, respectively, was identified, and an additional *ori* region unique to pIMP76_1 was detected, spanning positions 122,075– 122,503. Replicons of pIMP47 perfectly matched the IncHI1A(pNDM-CIT)/IncHI1B(pNDM-CIT)^1^ haplotype within the IncHI1AB subgroup of IncHI1 plasmids [13]. This haplotype was originally defined for plasmid pNDM-CIT, which was described in *Citrobacter freundii* [66] and carried genes conferring MDR (carbapenems, aminoglycosides, sulphonamides, trimethoprim, macrolides, phenicols, and bleomycin) as well as metal resistance (arsenic and tellurium). Plasmid pIMP76_1 also perfectly matched replicon types IncHI1A(pNDM-CIT) and IncHI1B(pNDM-CIT) and closely matched IncR, differing by two nucleotide substitutions. The IncR replicon was originally defined for the 98-kbp IncR/IncFIA(HI1)/repB(R1701) plasmid pK245 (GenBank accession: DQ449578.1), previously described in *K. pneumoniae* [67] and found to confer MDR (β-lactams, aminoglycosides, sulphonamides, trimethoprim, tetracyclines, quinolones, and phenicols) (Figure 1). Plasmid pIMP76_2, which was predicted to be non-mobilisable due to the absence of any *oriT* site or genes coding for relaxases or the MPF, matched the IncFIB(K)(pCAV1099-114)/RepB/repFIB haplotype with 98–99.6% nucleotide identities and 100% coverage to replicon probes in the PlasmidFinder database.

**Figure 1.**
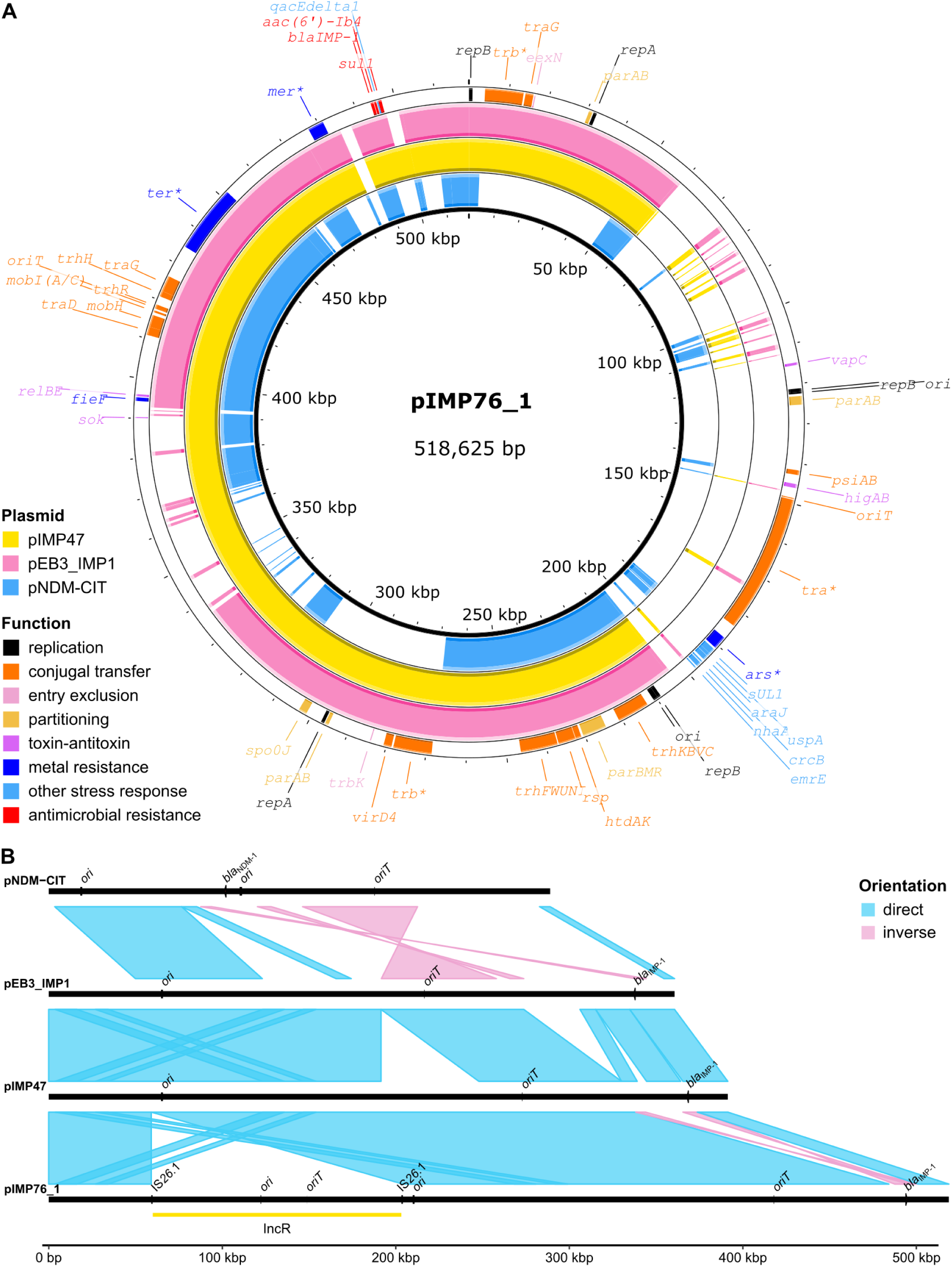
Sequence comparison between IncHI1A(NDM-CIT)/IncHI1B(pNDM-CIT) plasmids pIMP47, pIMP76_1, pEB3_IMP1 (reorientated), and pNDM-CIT (reorientated). Labels *ori* and *oriT* stand for the origin of replication and the origin of transfer, respectively. See Figures S1–3 in Supplementary Results for genetic structure and functions of pIMP47, pIMP76_1, and pEB3_IMP1. (A) Alignment of pIMP76_1 (query sequence) against pIMP47, pEB3_IMP1, and pNDM-CIT (subject sequences) using BLASTn as implemented in Proksee. Asterisks indicate large gene clusters. For example, *mer*\* represents the *merRTPCADE* cluster encoding mercury resistance. (B) Pairwise comparison between these four plasmids using megaBLAST, with aligned regions (>5 kbp) indicated by shades that represent a minimum nucleotide identity of 80% and are coloured by the orientations of alignments. The yellow bar denotes the 143-kbp exogenous IncR region flanked by IS*2c*.1 in pIMP76_1.

Both pIMP47 and pIMP76_1 carried genes conferring multidrug resistance (3–6 antimicrobial classes) and resistance to mercury (*merRTPCADE* genes) and tellurium (*terYXWZABCDEF* genes) [68, 69]. Moreover, pIMP47 harboured a *chrA* gene conferring chromium resistance [70] and a *qacE* gene (Table 2) conferring resistance to quaternary ammonium compounds, biocides that have been widely used since the 1930s [71]. By contrast, pIMP76_2, which did not carry any AMR gene, was enriched with genes conferring resistance to copper (*pco* genes), silver (*sil* genes), arsenic (*ars* genes), and heat (*hsp20*, *clpK*, *shsP*, *yfdX1, yfdX2, hdeD-GI*, *trxLHR*, *kefB-GI*, and *psi-GI*).

### Homologues of pIMP47, pIMP76_1, and pIMP76_2 in GenBank

Plasmid pIMP76_1 possessed a unique 143,354-bp IncR region (positions 59,901–203,254) bounded by two directly-orientated copies of a 100A>G variant of IS*26* (hereafter, IS*2c*.1) in the reverse complementary strand. No other copy of IS*2c*.1 was identified in pIMP76_1. Notably, the IS*2c*.1–IncR– IS*2c*.1 structure (144,994 bp; positions 59,081– 204,074) in pIMP76_1 was flanked by 8-bp direct repeats (DRs) of CCCAAAAG in the reverse complementary strand, and no DRs were found to flank each individual IS*2c*.1. This genetic structure and DR pattern suggest IS*26*-mediated cointegration or fusion between an exogenous IncR segment, as either a translocatable unit or pseudo composite transposon, and an IncHI1 plasmid, although the exact pathway could not be determined [72–74]. The IncR region exhibited the closest match with the 100,945-bp IncR plasmid pKPNB_13380.3 (CP154206.1) from *K. pneumoniae*, with an overall >99% nucleotide identity over 67% of this region. By contrast, only 7.7 kbp of this IncR region matched with plasmid pK245 (positions 13,961–21,684, with 97% nucleotide identity) besides insertion sequences, and the aligned region comprised partition genes *parAB* and the replication gene *repB*.

Only partial matches with plasmids pIMP47 and pIMP76_1 were identified in GenBank, while near identical plasmids were found for pIMP76_2. Notably, a 361-kbp *bla*_IMP-1_-carrying IncHI1A(pNDM-CIT)/IncHI1B(pNDM-CIT) plasmid pEB3_IMP1 (CP141538.1; Figure S3, Supplementary Results), originally identified in an *Enterobacter hormaechei* isolate recovered from a nosocomial faecal sample in South West England in 2022 [75], shared >99% nucleotide identity across 86% of pIMP47 and 63% of pIMP76_1, respectively (Figure 1A). Annotation of pEB3_IMP1 with bakta revealed that this plasmid also carried the *merRTPCADE* and *terYXWZABCDEF* gene clusters. Plasmid pIMP76_2 shared 99% nucleotide identity over 99% of its length with the 209-kbp IncFIB(K)/RepB/repFIB plasmids pNK_H5_007.1 (CP153418.1, from *K. pneumoniae*) and pKGVET2019-01-792.1 (CP154454.1, from *K. variicola*), recovered in Norway in 2017 and 2019, respectively.

### Genetic context of *bla*_IMP-1_ across IncHI1 plasmids

We resolved genetic structures encompassing *bla*_IMP-1_ in plasmids pEB3_IMP1, pIMP47, and pIMP76_1. An intact class 1 integron was found to carry both *bla*_IMP-1_ and *aac(6’)-Ib4* as gene cassettes in pIMP76_1 (Figure 2). This integron was 4,779 bp in length (positions 17,127– 21,905 in pIMP76_1) from the 5’-conserved segment (CS) to the 3’-CS [76]. It was also identified in pIMP47 and pEB3_IMP1 despite a 323-bp truncation resulting in a 178-bp remnant of *orf5* (*yhbs*) in the integron’s 3’-CS. These integrons were located in a highly dynamic region across these three plasmids (Figures 1B and 2). Screening the intact integron against the TnCentral database identified the *bla*_IMP-1_-harbouring integron In31 (GenBank accession: AJ223604.1), which was discovered in *Pseudomonas aeruginosa* and harbours an additional *catBc* cassette [76], as the closest match (99% nucleotide identity and 100% query coverage). Notably, homologues (96–99% nucleotide identity and 93–100% query coverage) of the *bla*_IMP-1_-harbouring integron from pIMP76_1 were detected in diverse plasmids across *Enterobacterales* species. Such plasmids included replicon types IncHI2 (e.g., CP044215.1 and CP043767.1) and IncN3 (e.g., CP043856.1), as previously reported in the UK [77], and a *P. aeruginosa* plasmid (CP096823.1) of an unknown type collected in China in 2015. Similar integrons were also found in the chromosome of *P. aeruginosa*, including those of strains IMP-13 (CP034354.1) and PA5957 (CP173023.1) from Belgium (2011) and France (2015), respectively.

**Figure 2.**
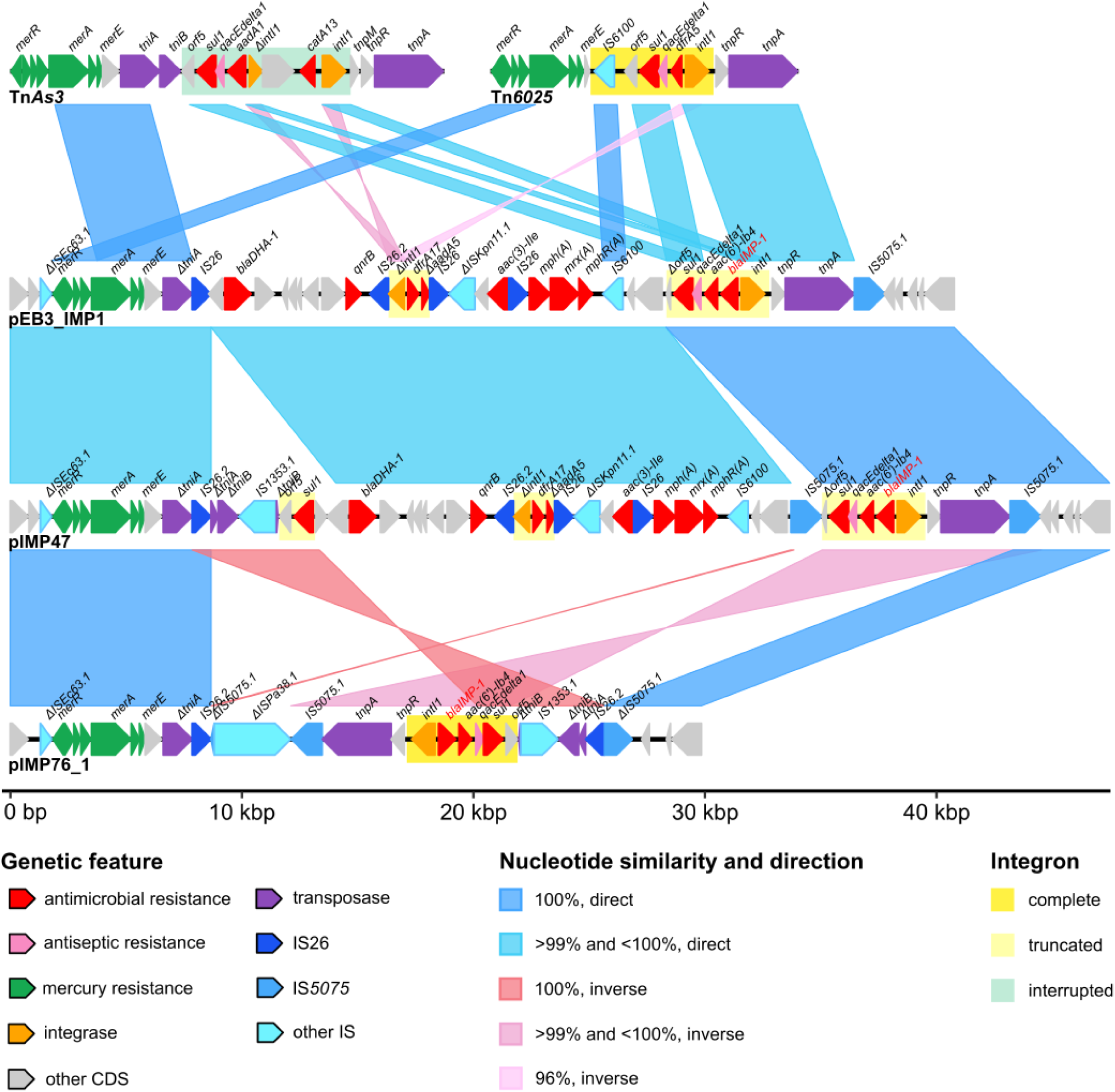
Comparison between the genetic context of *bla*_IMP-1_ in plasmids pIMP47, pIMP76_1, and pEB3_IMP1 and putative prototype transposons (Tn*As3* and Tn*c025*) containing class 1 integrons. To account for arbitrary deposition orientations and prevent artificial inversions, the reference sequence of pEB3_IMP1 (GenBank accession: CP141538) was reoriented with dnaapler v1.2.0, and the reverse complementary strand of Tn*c025*’s reference sequence (accession: GU562437.2) was used for this comparison. All sequences were annotated with bakta v1.11.3 and the full database v6.0 for consistency. The label of *bla*_IMP-1_ is highlighted in red, and integrons are indicated with yellow and green boxes in the background. The orientation of each insertion sequence (IS) was determined as per its reference sequence in the ISfinder database. Each unique IS variant is denoted with a decimal in the locus label, such as IS*2c*.2. See Tables S3–7 for sequence annotations. The abbreviation CDS stands for a coding sequence.

Flanking regions of the *bla*_IMP-1_-harbouring integrons in pEB3_IMP1, pIMP47, and pIMP76_1 exhibited high sequence homology (>99% overall nucleotide identity and 31–38% regional coverage) with the Tn*3*-family transposon Tn*c025* (positions 888–14,189 in GU562437.2) [78], which carries a class 1 integron (Figure 2). In Tn*c025*, a 5,003-bp segment (positions 22–5,024) comprising the transposase (TnpA) and resolvase (TnpR) genes, together with an integron 5’-CS, matched a 5,019-bp right-flanking region of *bla*_IMP-1_ in pEB3_IMP1, pIMP47, and pIMP76_1, exhibiting 99.6% nucleotide identity and differing by only two nucleotide substitutions and a 16-bp indel; and a 1,003-bp segment (positions 7,829–8,831) comprising an intact IS*c100* (IS*c*/IS*2c* family) was shared by pEB3_IMP1 and pIMP47 in the left flanking region of *bla*_IMP-1_, creating the same IS*c100*–integron–*tnpR*–*tnpA* structure across Tn*c025* and both plasmids.

### Association of *bla*_IMP-1_ with insertion sequences in IncHI1 plasmids

IS*c*/IS*2c*-family element IS*26* and its three variants (Supplementary Results, Pages 4–5), IS*2c*.1, IS*2c*.2 (702C>A), and IS*2c*.3 (614G>A and 736A>T), were identified in pEB3_IMP1, pIMP47, and pIMP76_1, with a total copy number of five in pEB3_IMP1, seven in pIMP47, and nine in pIMP76_1 (Table S8). In these plasmids, IS*2c* and IS*2c*.2 occurred within flanking regions of the *bla*_IMP-1_-harbouring integrons. Notably, a 7,370-bp ΔIS*Ecc3*.1–*merRTPCADE*–Δ*tniA*–IS*2c/*IS*2c.*2 segment upstream of these integrons was conserved across all three plasmids (Figure 2), with a single position differing between IS*2c* and IS*2c*.2. The ΔIS*Ecc3*.1 locus was a 3’-truncated IS*Ecc3* variant (positions 1–501 out of 4,473 bp remained), and Δ*tniA* was an IS*2c*/IS*2c*.2-interrupted transposase gene *tniA* of a Tn*As3*-like transposon in the Tn*3* family.

Within this 7.4-kbp segment, a 6,046-bp region comprising *merRTPCADE* and Δ*tniA* could be split into two parts based on homology. Specifically, positions 1–1,940 of this region precisely matched (100% nucleotide identity and query coverage) with Tn*c025* from *merR* to position 232 in *merA*; and positions 1,941–6,046 precisely matched with Tn*As3* from position 233 in *merA* to position 1,252 in *tniA* (Figure 2). Such a split suggests insertion of a Tn*c025*-like transposon at an IS*Ecc3*-like element in a precursor of the three plasmids sequentially followed by recombination between the precursor and the Tn*As3*-like transposon within this 6-kbp region and insertion of an IS*2c* or IS*2c*.2 that interrupted the *tniA* gene.

The *bla*_IMP-1_-harbouring integron was also associated with the IS*110*-family element IS*5075* in these plasmids, which only harboured a novel variant (designated as IS*5075*.1) exhibiting four single-nucleotide substitutions (Supplementary Results, Pages 6–7). This variant was detected in genomes of *P. aeruginosa*, *P. marginalis*, *Acinetobacter baumannii*, and across *Enterobacterales* species in GenBank using megaBLAST. IS*5075*.1 had smaller copy numbers than IS*26*-like elements: one in pEB3_IMP1, two in pIMP47, and one intact copy plus two truncation remnants (positions 1–134 and 127–1,327; truncated by two copies of IS*2c*.2) in pIMP76_1. All six full or partial matches were located in the integron’s flanking regions. Notably, a copy of IS*5075*.1 was located downstream of an intact copy of Tn*c025*’s transposase gene *tnpA* in all three plasmids, forming a 9–10 kbp conserved ‘*bla*_IMP-1_-harbouring integron–*tnpR*–*tnpA*– IS*5075*.1’ segment (9,769 bp in pIMP76_1 and 9,446 bp in pEB3_IMP1 and pIMP47) including the 3’-CS of the integron (Figure 2). In pIMP76_1, which harboured the simplest genetic structure of *bla*_IMP-1_, the two IS*5075.1* remnants flanking the integron in inverse directions and truncation of an IS*Pa38*-like element on both sides (Figure 2 and Table S7) implicate a putative IS*5075*.1-mediated inversion of the bounded region via recombination after the truncation of the IS*Pa38*-like element and before both IS*5075*.1 elements at the boundaries got truncated by IS*2c*.2.

In pIMP47, two directly-oriented copies of IS*5075*.1 flanking the *bla*_IMP-1_-harbouring integron as well as the Tn*c025*-like resolvase (TnpR) and transposase (TnpA) genes *tnpRA* formed a 10,794-bp hypothetical composite transposon (Figure 2). This direct orientation contrasts with the inverse orientation of IS*5075* elements observed in composite transposon Tn*5075* (GenBank accession: AF457211) [79]. Search of this hypothetical transposon in the NCBI nucleotide databases for bacteria and plasmids using megaBLAST and in the PLSDB database v2024_05_31_v2 using both the MASH-screen and MASH-dist methods under default parameters did not identify any hit of a 100% query coverage, indicating a lack of evidence to support the hypothesised mobility. Similarly, no evidence of transposon-mediated mobility was found using the same search methods for a 17,808-bp segment bounded by two inversely-orientated copies of IS*2c*.2 in the integron’s flanking regions in pIMP76_1 (Figure 2), which is consistent with the cointegration mechanism of IS*26*’s activity [80].

### Comparative analysis of conjugation machinery

Both pIMP47 and pIMP76_1 were predicted to encode proteins for the type-F MPF and shared the same *oriT* site spanning positions 272,671–272,954 in pIMP47 and 417,673–417,956 in pIMP76_1, respectively. Both plasmids encoded relaxases in the MOBH and MOBP families. Moreover, the unique IncR region of pIMP76_1 possessed a unique *oriT* site spanning positions 148,684–148,733 (Figure 1) as well as a *traI* gene encoding a MOBF-family relaxase (Figure 3).

**Figure 3.**
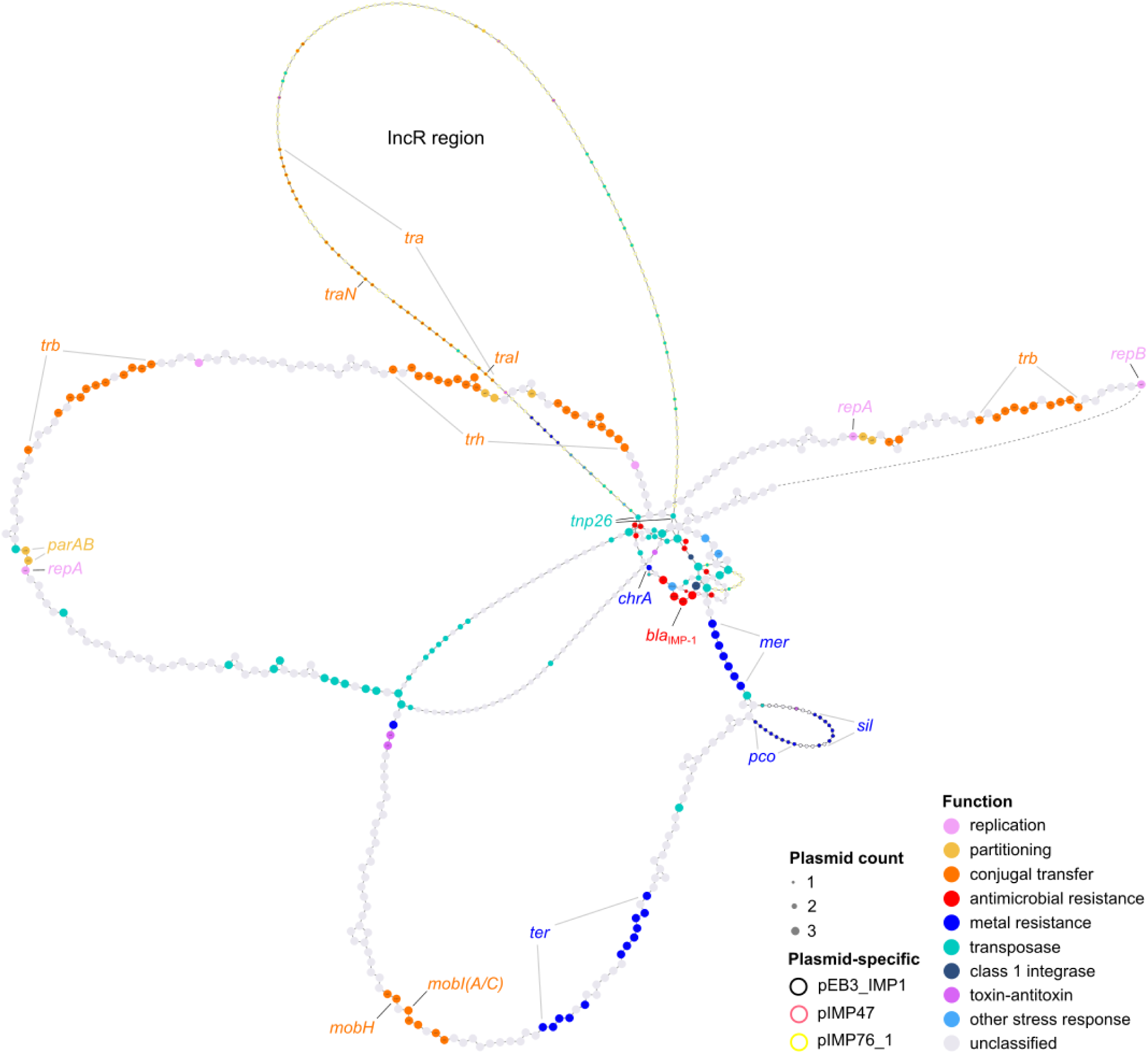
Pangenome graph of plasmids pIMP47, pIMP76_1, and pEB3_IMP1. Each node represents a cluster of orthologous genes (COG, also known as a gene family) determined with Panaroo based on an identity threshold of 70% for protein sequences and genetic context, and each edge indicates physical adjacency between two COGs in plasmid sequences. The node diameter represents the number of plasmids carrying a specific COG, and the border colour, if present, denotes a plasmid-specific COG. Paired grey lines indicate gene clusters, and black lines indicate individual genes. The dashed edge connects the arbitrary start (*repB*) and end (an intergenic region downstream of a hypothetical gene) of all three plasmid sequences. This edge is missing from the original pangenome graph as a technical artefact because Panaroo did not account for the topology of input circular genomes. The *tnp2c* gene encodes the transposase of IS*2c*.

The *tra*, *trb*, *trh* and *htd* genes encoding conjugation systems were compared among pIMP47, pIMP76_1, and pEB3_IMP1 to evaluate the conjugative potential of these plasmids. Pangenome analysis of clusters of orthologous genes (COGs) revealed that all three plasmids shared two *trbBCDEFIJ* clusters, one *trhBCFIKNUVW* cluster, one *htdAKOT* cluster, and the individual genes *traD*, *traG*, *traF*, *trbK*, *trhH*, and *trhR*, whereas a *traABCDEFGHIKLMNQSTUVWX* cluster was identified only within pIMP76_1’s IncR region (Figure 3).

Robust global alignments of TraN/TrhN sequences were obtained only within each type, and phylogenetic trees were reconstructed for the short and long types, respectively (Figures 4A and 4B). TraN_S1_, a 651-aa protein encoded by pIMP76_1’s unique *traN* gene, differed from TraN_S_β1 in the β subtype of the short TraN by five amino acid substitutions (R29S, T70A, S196N, N261S, S345N), including two located in the shared tip domain T177-V335 (Figure 4C) [54]. This subtype is specialist associated with species-specific conjugation [81]. The *trhN* gene in pIMP47 and pEB3_IMP1 encoded the same protein TrhN_1_, which differed from that (TrhN_2_) encoded by pIMP76_1 by a single amino acid substitution E2K.

**Figure 4.**
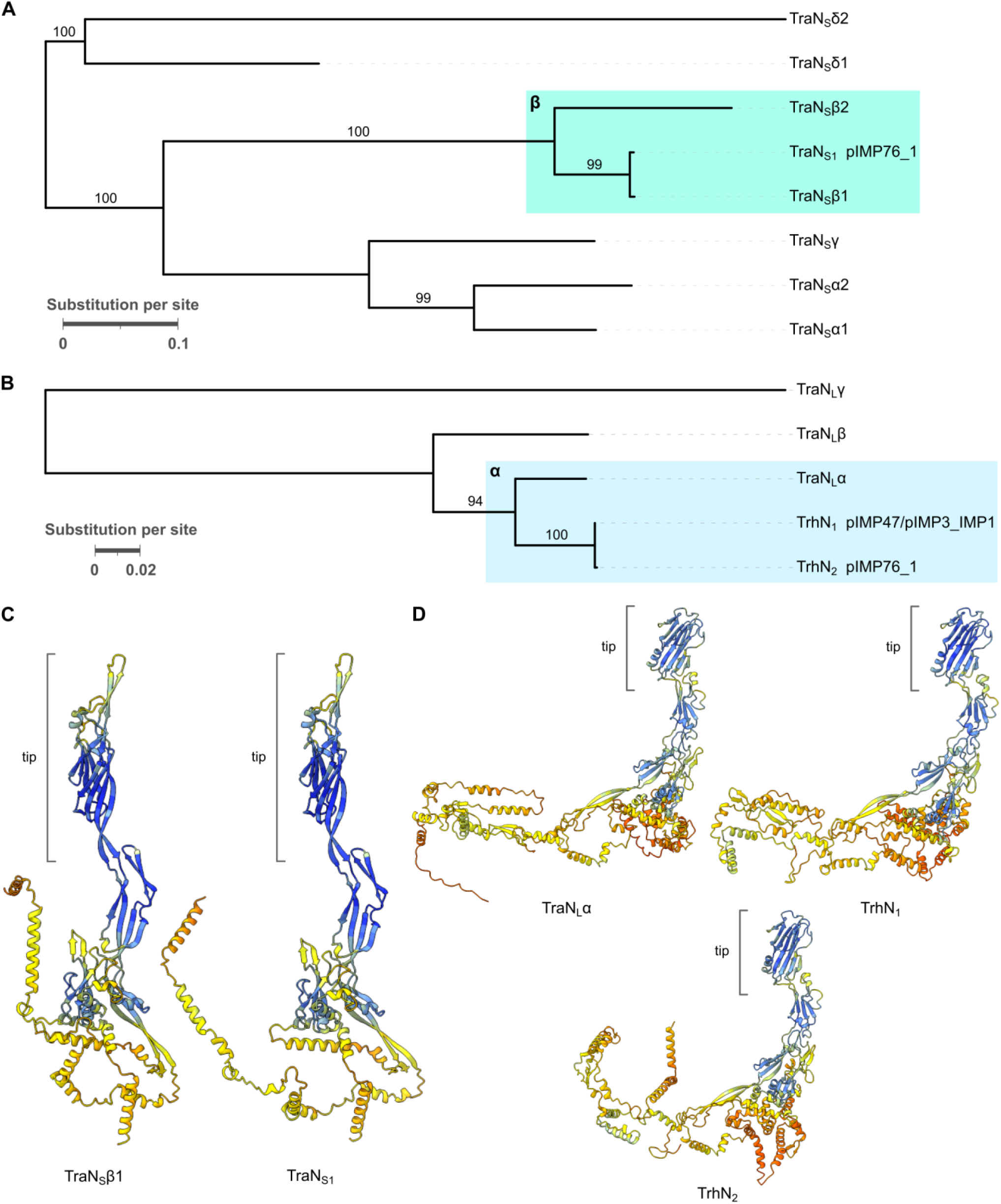
Midpoint-rooted maximum-likelihood phylogenetic trees of short (S) and long (L) TraN sequences (A, B) and predicted three-dimensional structures of selected short and long TraN sequences (C, D), with tip domains indicated by the grey bars. Bootstrap values are noted above internal branches in the trees. The β subtype of the short TraN and α subtype of the long TraN are labelled and denoted by the green and blue shades, respectively. Amino acid chains in the structural models are coloured by AlphaFold3’s pLDDT confidence scores using ChimeraX’s alphafold palette.

Both TrhN_1_ and TrhN_2_ belonged to the α subgroup of the long TraN, differing from the representative sequence TraN_L_α, originally identified in the IncHI1 plasmid R27 [55], by 66 and 67 amino acids (94% amino acid identity), respectively. Nevertheless, TrhN_1_, TrhN_2_, and TraN_L_α shared the same tip domain S305-E457 (Figure 4D) [55].

### Experimental assessment of plasmid mobility

Putative transconjugants were obtained from the mating mixture of IMP76 and ICC8001 at an end-point OD of 0.149, with an average of 99 colonies per rifampicin–ertapenem plate (range: 79–115). Three randomly selected transconjugant colonies exhibited the same level of rifampicin MICs (>512 mg/L) as ICC8001 and ertapenem MICs (>2 mg/L) as IMP76. Both IMP47 and IMP76 were negative in the string test (string lengths <1 mm), whereas ICC8001 and the three selected transconjugants were hypermucoviscous. By contrast, no transconjugant was obtained from the mating mixture of IMP47 and ICC8001 at an end-point OD of 0.166 (see Table S9 for ODs of all cultures). As expected, lawns of IMP47, IMP76, and both mating mixtures (IMP47-ICC8001, IMP76-ICC8001) grew on ertapenem-only and antimicrobial-free plates; and lawns of ICC8001 and the mating mixtures grew on rifampicin-only and antimicrobial-free plates, suggesting that rifampicin and ertapenem concentrations in the rifampicin– ertapenem plates were adequate to inhibit the growth of non-transconjugants in both mixtures. Collectively, these results confirm the conjugative ability of pIMP76_1, whereas conjugative transfer was not observed for pIMP47 under the tested conditions.

## Discussion

In this study, we have described complete genomes and plasmids of *K. grimontii* isolate IMP47 and *K. pneumoniae* isolate IMP76 identified in two inpatients during routine screening of CPE carriage. Both isolates were associated with hospital stays in the same geographical area. However, time and source of gut colonisation could not be inferred for either patient owing to a lack of prior screening. The *bla*_IMP-1_-carrying IncHI1 megaplasmids, pIMP47 (391 kbp) and pIMP76_1 (519 kbp), were unique in GenBank and carried diverse genes coding for MDR and stress responses. We identified pEB3_IMP1, another *bla*_IMP-1_-carrying IncHI1 plasmid in GenBank, as the closest match to both pIMP47 and pIMP76_1. Comparison between these three plasmids elucidated that *bla*_IMP-1_ was harboured in nearly identical class 1 integrons with flanking regions sharing 5–10 kbp conserved segments related to Tn*As3* and Tn*c025*, explaining the physical linkage between *bla*_IMP-1_ and the *merRTPCADE* cluster in these plasmids. Such comparative analysis also revealed the association between TEs (ISs and transposons) and *bla*_IMP-1_ in these plasmids. Globally, *bla*_IMP-1_ was one of the most frequent alleles of *bla*_IMP_ as of 2023 [12] and has spread across species of *Pseudomonas*, *Acinetobacter*, and *Enterobacterales* [8], being detected in regional outbreaks particularly in hospitals [19, 82–84], where sinks in patient wards have been implicated as a persistent reservoir [85–87]. The highly homologous segments shared between Tn*As3*, Tn*c025*, and flanking regions of the *bla*_IMP-1_-harbouring class 1 integron in plasmids pEB3_IMP1, pIMP47, and pIMP76_1 (Figure 2) suggest a single acquisition event of a Tn*c025*-like transposon in an ancestral plasmid followed by recombination with a Tn*As3*-like transposon from position 233 in *merA*. The order of insertion events can be inferred for ISs in short plasmid segments based on truncation or interruption patterns (Figure 2 and Tables S5–7). The diverse complete or partial ISs and Tn*As3/6025*-like segments in flanking regions of the *bla*_IMP-1_-harbouring integron in the three investigated plasmids may facilitate further recombination events, resulting in transposition or horizontal transfer of this integron.

IS*2c* is a well-characterised driver of AMR dissemination among Gram-negative bacteria [88]. IS*2c* and/or its variants, which collectively had 5–9 copies in pIMP47, pIMP76_1, and pEB3_IMP1, may mediate the mobilisation of *bla*_IMP-1_ in three pathways. First, such an IS may excise with its adjacent, *bla*_IMP-1_-harbouring sequence region in a plasmid through replicative transposition of the IS and form a translocatable unit, which can be incorporated at a distinct genomic location [72]. Second, when bounded by directly-oriented IS*c*/IS*2c*-family elements, the *bla*_IMP-1_-harbouring integron can transpose as a pseudo composite transposon [74]. Third, IS*26*-mediated transposition may introduce an additional conjugation system to a *bla*_IMP-1_-carrying plasmid, as exemplified by this study, where pIMP76_1 uniquely acquired *tra* genes and an *oriT* site in the exogeneous IncR region, as well as established cases of IS*2c*-mediated plasmid cointegration [89, 90].

Both genomic and molecular analyses in literature have revealed that IncH plasmids have a broad host range, circulating specifically among *Enterobacterales* across One Health sectors [13, 55]. This is in line with the taxonomical range of plasmid hosts where long TraN proteins (including TrhN_1_ and TrhN_2_ reported in this study) were isolated [55]. Nonetheless, pIMP47 and pIMP76_1 exhibited distinct conjugative capacity in our study, where the predicted conjugation could only be experimentally validated for pIMP76_1 under the same condition despite the shared IncHI1 conjugation machinery (*oriT*, MOBH and MOBP relaxases, and the *trb*, *trh*, and *htd* gene clusters). The successful conjugal transfer of pIMP76_1 in the mating experiment could be attributed to the acquired conjugation machinery (*tra* genes and the unique *oriT* site) in the IncR region of this plasmid or the identity between the donor and recipient species comprising the mating mixtures (IMP76-ICC8001: same species; IMP47-ICC8001: two species complexes) or both. Further experiments are needed to evaluate the conjugative potential of pIMP47 and to determine the conjugation frequency and fitness cost of pIMP76_1.

The co-carriage of *bla*_IMP-1_ and metal-resistance genes such as the *mer* and *ter* clusters in pIMP47, pIMP76_1, and pEB3_IMP1 (Figure 3) potentially provides these megaplasmids with an evolutionary advantage by facilitating their co-selection and maintenance in bacterial communities upon exposure to environmental stressors. Megaplasmids with this genetic configuration, such as three other *bla*_IMP-1_-carrying IncHI1A(pNDM-CIT)/IncHI1B(pNDM-CIT) plasmids previously reported in two regions in the UK (sequences are not publicly available) [75], may act as flexible scaffolds for disseminating AMR determinants in the healthcare environment, given the implicated role of hospital wastewater systems as a persistent reservoir for CPE [91].

Taken together, our study suggests monitoring IncHI1 plasmids as a putative emerging vector of *bla*_IMP-1_ and its derivatives, emphasising tracking dynamics of *bla*_IMP-1_’s associated genetic structures for understanding the spread of this resistance determinant. Further studies are warranted to determine conjugation frequencies and fitness costs of *bla*_IMP-1_-carrying plasmids across diverse environmental temperatures, chemical stressors, and bacterial hosts. Longitudinal investigation of TE-mediated genomic rearrangement and horizontal gene transfer will be crucial for predicting the emergence of new MDR variants. Ultimately, understanding how AMR-conferring megaplasmids interact with healthcare-associated microbiota, particularly between patients and the hospital environment, is essential for developing targeted interventions to disrupt the spread of high-risk resistance genes such as *bla*_IMP_.

## Conclusions

We have elucidated genetic features in two novel *bla*_IMP-1_-carrying IncHI1 megaplasmids from two gut-colonising CPE isolates. Through analysis of TEs, we show associations between AMR and metal resistance—particularly, between transferable carbapenem resistance and mercury resistance—and highlight the potential for further mobilisation of the *bla*_IMP-1_-carrying class 1 integron via TE-mediated mechanisms, which should be monitored via longitudinal genomic surveillance.

## Ethical statement

This study was conducted in accordance with the institutional Ethics Reference 21/LO/0170 (279677) and Protocol 21HH6538 ‘Investigation of Epidemiological and Pathogenic Factors Associated with Infectious Diseases’ at Imperial College London.

## Author contributions

**Yu Wan**: Conceptualisation, Methodology, Investigation, Data curation, Formal analysis, Visualisation, Writing – original draft, Writing – review and editing, Supervision, Project administration, Funding acquisition. **Victoria Orr**: Methodology, Investigation, Formal analysis, Visualisation, Writing – original draft, Writing – review and editing. **Maria Getino**: Conceptualisation, Methodology, Investigation, Data curation, Formal analysis, Writing – review and editing. **Sophie Mannix**: Formal analysis, Writing - original draft, Writing – review and editing. **Chloe Heenan**, **Ebony Richmond-Mensah**, and **Joshua L. C. Wong**: Investigation. **Rojus Urbonas**, **Martina O. Chukwu**, and **Jane F. Turton**: Investigation, Writing – review and editing. **Nicholas Harper**: Methodology. **Katie L. Hopkins** and **Gad Frankel**: Resources, Writing – review and editing. **Alison H. Holmes**: Funding acquisition. **Frances Davies**: Resources, Funding acquisition, Writing – review and editing. **Elita Jauneikaite**: Conceptualisation, Methodology, Resources, Investigation, Data curation, Supervision, Project administration, Funding acquisition, Writing – review and editing.

## Supporting information

Supplementary results

## Acknowledgment

We acknowledge the Colebrook Laboratory, a facility supported by the NIHR Imperial Biomedical Research Centre (BRC), for providing resources for microbiology experiments, nanopore sequencing, and bioinformatics analysis. Part of the bioinformatics analysis was performed on equipment purchased as part of MRC CARP fellowship award MR/T005254/1. We also acknowledge use of the High-performance Computing platform provided by Liverpool Shared Research Facilities in the Faculty of Health and Life Sciences at the University of Liverpool. We express our gratitude to the Antimicrobial Pharmacodynamics and Therapeutics (APT) group at the University of Liverpool for sharing equipment and consumables and thank Dr Nada Reza, Katarzyna Jezewska, Dr Vineet Dubey, Charlotte M. Jones, Iona Horner, and Dr Nicola Farrington in the APT group for sharing laboratory experience. We thank Michael Gilmore and Anna Morkowska at Imperial College Healthcare NHS Trust and Julia Sanchez-Garrido at Imperial College London for facilitating isolate transfer. We are grateful to the Institut Pasteur teams for the curation and maintenance of BIGSdb-Pasteur databases at bigsdb.pasteur.fr.

## Footnotes

1 The IncHI1A(pNDM-CIT) replicon is named as IncHI1A(NDM-CIT) in the PlasmidFinder database as of the latest version (v2025-11-27) accessed in this study.

