## Supplementary results for "Two IncHI1 megaplasmids in *Klebsiella* species reveal transposable-element-mediated *bla*IMP-1 mobilisation"

Yu Wan, Victoria Orr, Maria Getino, Sophie Mannix, Chloe Heenan, Ebony Richmond-Mensah, Nicholas Harper, Joshua L. C. Wong, Rojus Urbonas, Martina O. Chukwu, Jane F. Turton, Katie L. Hopkins, Gad Frankel, Alison H. Holmes, Frances Davies, and Elita Jauneikaite

September 2026

**Figure S1**

Structure and functions of genetic features in plasmid pIMP47. The grey histogram in the inner ring illustrates the GC content calculated by Proksee using a 1-kbp sliding window and 10-bp steps.

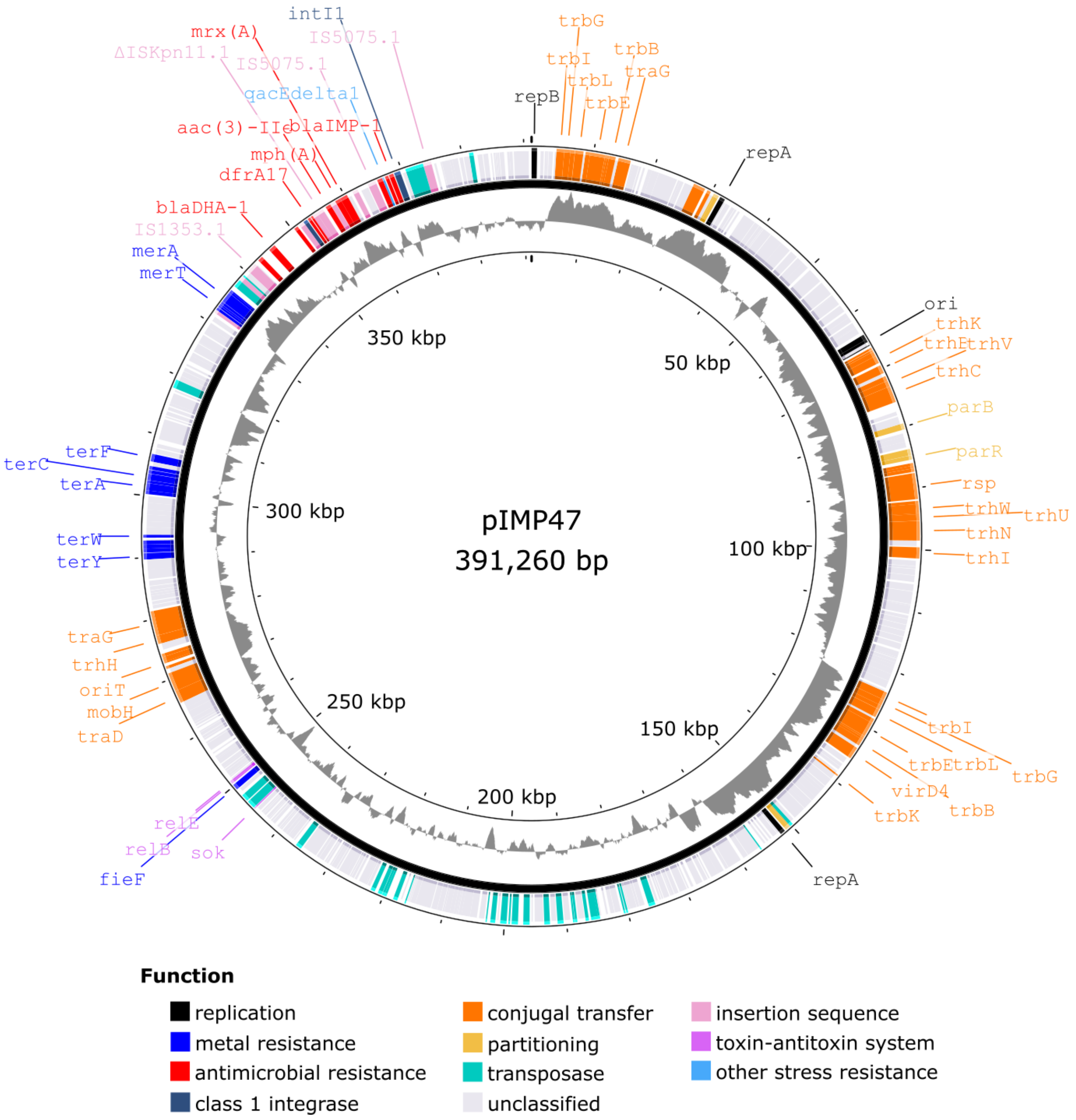

### Figure S2

Structure and functions of genetic features in plasmid pIMP76_1. The grey histogram in the inner ring illustrates the GC content calculated by Proksee using a 1-kbp sliding window and 10-bp steps.

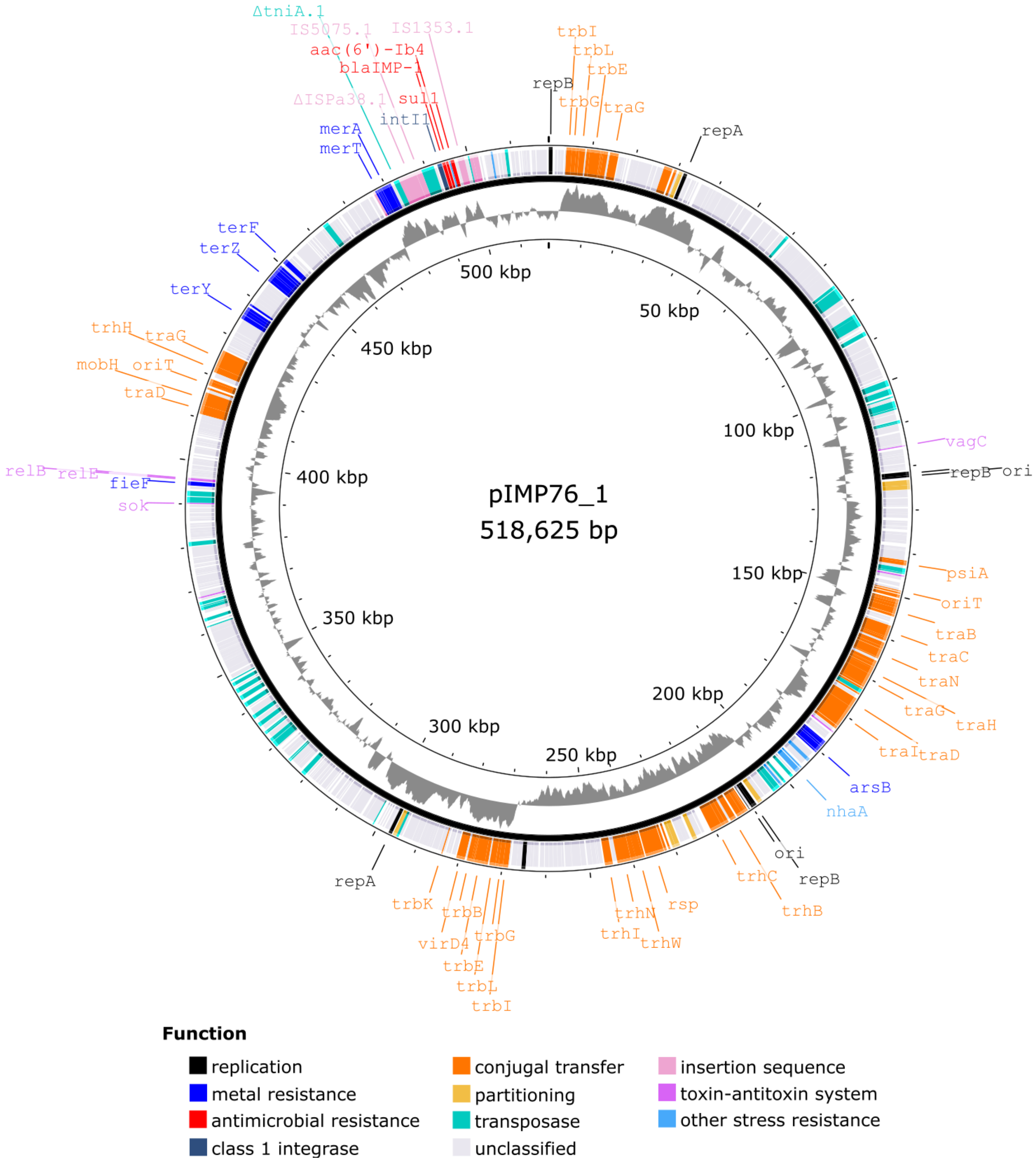

### Figure S3

Structure and functions of genetic features in plasmid pEB3_IMP1. The grey histogram in the inner ring illustrates the GC content calculated by Proksee using a 1-kbp sliding window and 10-bp steps.

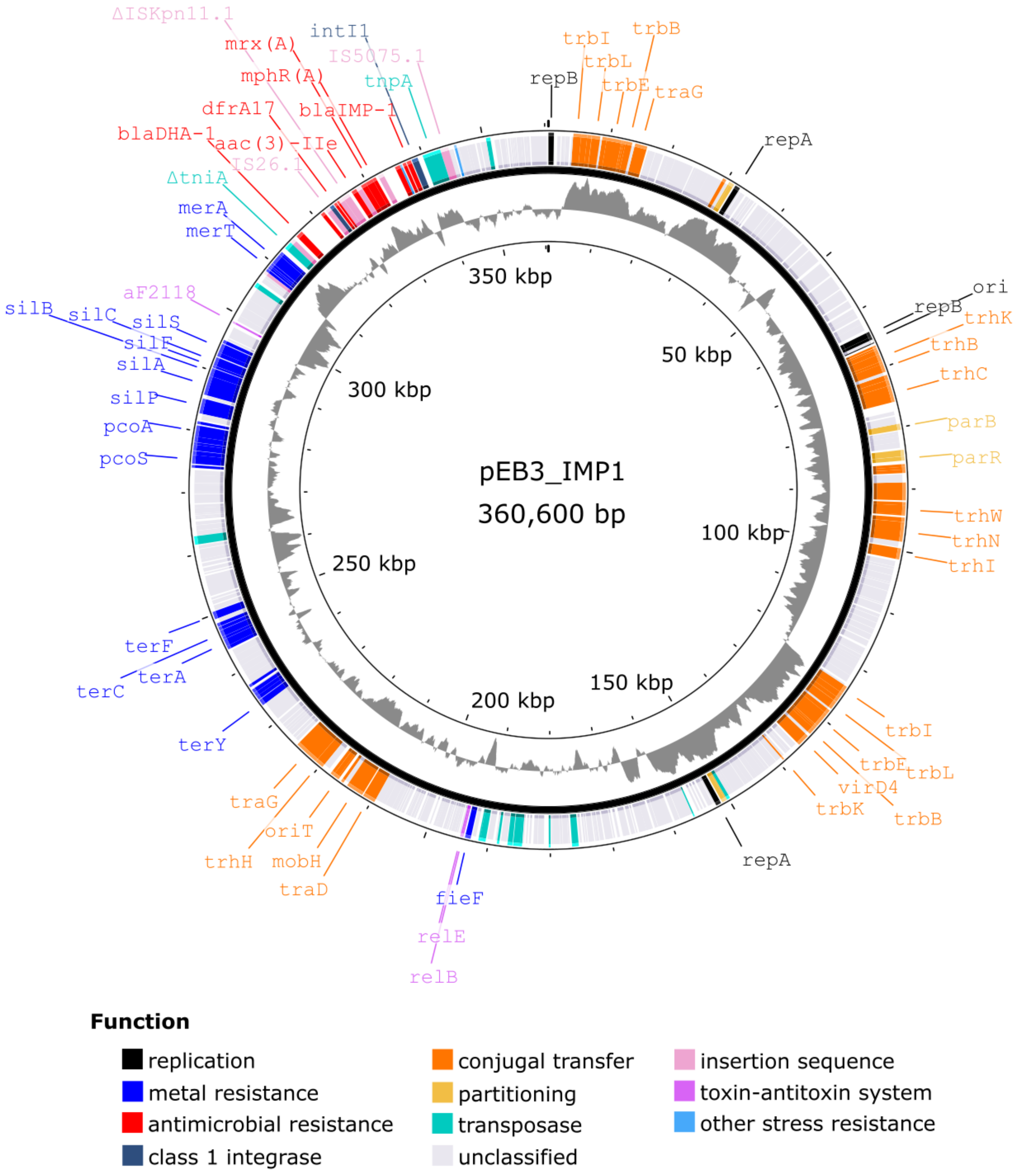

### Multisequence alignment of IS*26* and variants

Sequences of IS*26* and three variants (IS*26*.1–3) identified in IncHI1 plasmids pEB3_IMP1, pIMP47, and pIMP76_1 were aligned using Clustal Omega v1.2.4 implemented on the European Bioinformatics Institute (EMBL-EBI) website (www.ebi.ac.uk/jdispatcher/msa/clustalo). The alignment is presented in the CLUSTAL format, where an asterisk beneath the alignment indicates an invariable position.

```
IS26.3      GGCACTGTTGCAAATAGTCGGTGGTGATAAACTTATCATCCCCTTTTGCTGATGGAGCTG      60
IS26.1      GGCACTGTTGCAAATAGTCGGTGGTGATAAACTTATCATCCCCTTTTGCTGATGGAGCTG      60
IS26        GGCACTGTTGCAAATAGTCGGTGGTGATAAACTTATCATCCCCTTTTGCTGATGGAGCTG      60
IS26.2      GGCACTGTTGCAAATAGTCGGTGGTGATAAACTTATCATCCCCTTTTGCTGATGGAGCTG      60
            ************************************************************

IS26.3      CACATGAACCCATTCAAAGGCCGGCATTTTCAGCGTGACATCATTCTGTGGGCCGTACGC      120
IS26.1      CACATGAACCCATTCAAAGGCCGGCATTTTCAGCGTGACGTCATTCTGTGGGCCGTACGC      120
IS26        CACATGAACCCATTCAAAGGCCGGCATTTTCAGCGTGACATCATTCTGTGGGCCGTACGC      120
IS26.2      CACATGAACCCATTCAAAGGCCGGCATTTTCAGCGTGACATCATTCTGTGGGCCGTACGC      120
            *************************************** ********************

IS26.3      TGGTACTGCAAATACGGCATCAGTTACCGTGAGCTGCAGGAGATGCTGGCTGAACGCGGA      180
IS26.1      TGGTACTGCAAATACGGCATCAGTTACCGTGAGCTGCAGGAGATGCTGGCTGAACGCGGA      180
IS26        TGGTACTGCAAATACGGCATCAGTTACCGTGAGCTGCAGGAGATGCTGGCTGAACGCGGA      180
IS26.2      TGGTACTGCAAATACGGCATCAGTTACCGTGAGCTGCAGGAGATGCTGGCTGAACGCGGA      180
            ************************************************************

IS26.3      GTGAATGTCGATCACTCCACGATTTACCGCTGGGTTCAGCGTTATGCGCCTGAAATGGAA      240
IS26.1      GTGAATGTCGATCACTCCACGATTTACCGCTGGGTTCAGCGTTATGCGCCTGAAATGGAA      240
IS26        GTGAATGTCGATCACTCCACGATTTACCGCTGGGTTCAGCGTTATGCGCCTGAAATGGAA      240
IS26.2      GTGAATGTCGATCACTCCACGATTTACCGCTGGGTTCAGCGTTATGCGCCTGAAATGGAA      240
            ************************************************************

IS26.3      AAACGGCTGCGCTGGTACTGGCGTAACCCTTCCGATCTTTGCCCGTGGCACATGGATGAA      300
IS26.1      AAACGGCTGCGCTGGTACTGGCGTAACCCTTCCGATCTTTGCCCGTGGCACATGGATGAA      300
IS26        AAACGGCTGCGCTGGTACTGGCGTAACCCTTCCGATCTTTGCCCGTGGCACATGGATGAA      300
IS26.2      AAACGGCTGCGCTGGTACTGGCGTAACCCTTCCGATCTTTGCCCGTGGCACATGGATGAA      300
            ************************************************************

IS26.3      ACCTACGTGAAGGTCAATGGCCGCTGGGCGTATCTGTACCGGGCCGTCGACAGCCGGGGC      360
IS26.1      ACCTACGTGAAGGTCAATGGCCGCTGGGCGTATCTGTACCGGGCCGTCGACAGCCGGGGC      360
IS26        ACCTACGTGAAGGTCAATGGCCGCTGGGCGTATCTGTACCGGGCCGTCGACAGCCGGGGC      360
IS26.2      ACCTACGTGAAGGTCAATGGCCGCTGGGCGTATCTGTACCGGGCCGTCGACAGCCGGGGC      360
            ************************************************************

IS26.3      CGCACTGTCGATTTTTATCTCTCCTCCCGTCGTAACAGCAAAGCTGCATACCGGTTTCTG      420
IS26.1      CGCACTGTCGATTTTTATCTCTCCTCCCGTCGTAACAGCAAAGCTGCATACCGGTTTCTG      420
IS26        CGCACTGTCGATTTTTATCTCTCCTCCCGTCGTAACAGCAAAGCTGCATACCGGTTTCTG      420
IS26.2      CGCACTGTCGATTTTTATCTCTCCTCCCGTCGTAACAGCAAAGCTGCATACCGGTTTCTG      420
            ************************************************************
```

```
IS26.3      GGTAAAATCCTCAACAACGTGAAGAAGTGGCAGATCCCGCGATTCATCAACACGGATAAA      480
IS26.1      GGTAAAATCCTCAACAACGTGAAGAAGTGGCAGATCCCGCGATTCATCAACACGGATAAA      480
IS26        GGTAAAATCCTCAACAACGTGAAGAAGTGGCAGATCCCGCGATTCATCAACACGGATAAA      480
IS26.2      GGTAAAATCCTCAACAACGTGAAGAAGTGGCAGATCCCGCGATTCATCAACACGGATAAA      480
            ************************************************************

IS26.3      GCGCCCGCCTATGGTCGCGCGCTTGCTCTGCTCAAACGCGAAGGCCGGTGCCCGTCTGAC      540
IS26.1      GCGCCCGCCTATGGTCGCGCGCTTGCTCTGCTCAAACGCGAAGGCCGGTGCCCGTCTGAC      540
IS26        GCGCCCGCCTATGGTCGCGCGCTTGCTCTGCTCAAACGCGAAGGCCGGTGCCCGTCTGAC      540
IS26.2      GCGCCCGCCTATGGTCGCGCGCTTGCTCTGCTCAAACGCGAAGGCCGGTGCCCGTCTGAC      540
            ************************************************************

IS26.3      GTTGAACACCGACAGATTAAGTACCGGAACAACGTGATTGAATGCGATCATGGCAAACTG      600
IS26.1      GTTGAACACCGACAGATTAAGTACCGGAACAACGTGATTGAATGCGATCATGGCAAACTG      600
IS26        GTTGAACACCGACAGATTAAGTACCGGAACAACGTGATTGAATGCGATCATGGCAAACTG      600
IS26.2      GTTGAACACCGACAGATTAAGTACCGGAACAACGTGATTGAATGCGATCATGGCAAACTG      600
            ************************************************************

IS26.3      AAACGGATAATCGACGCCACGCTGGGATTTAAATCCATGAAGACGGCTTACGCCACCATC      660
IS26.1      AAACGGATAATCGGCGCCACGCTGGGATTTAAATCCATGAAGACGGCTTACGCCACCATC      660
IS26        AAACGGATAATCGGCGCCACGCTGGGATTTAAATCCATGAAGACGGCTTACGCCACCATC      660
IS26.2      AAACGGATAATCGGCGCCACGCTGGGATTTAAATCCATGAAGACGGCTTACGCCACCATC      660
            ************* **********************************************

IS26.3      AAAGGTATTGAGGTGATGCGTGCACTACGCAAAGGCCAGGCCTCAGCATTTTATTATGGT      720
IS26.1      AAAGGTATTGAGGTGATGCGTGCACTACGCAAAGGCCAGGCCTCAGCATTTTATTATGGT      720
IS26        AAAGGTATTGAGGTGATGCGTGCACTACGCAAAGGCCAGGCCTCAGCATTTTATTATGGT      720
IS26.2      AAAGGTATTGAGGTGATGCGTGCACTACGCAAAGGCCAGGCATCAGCATTTTATTATGGT      720
            ***************************************** ******************

IS26.3      GATCCCCTGGGCGAATTGCGCCTGGTAAGCAGAGTTTTTGAAATGTAAGGCCTTTGAATA      780
IS26.1      GATCCCCTGGGCGAAATGCGCCTGGTAAGCAGAGTTTTTGAAATGTAAGGCCTTTGAATA      780
IS26        GATCCCCTGGGCGAAATGCGCCTGGTAAGCAGAGTTTTTGAAATGTAAGGCCTTTGAATA      780
IS26.2      GATCCCCTGGGCGAAATGCGCCTGGTAAGCAGAGTTTTTGAAATGTAAGGCCTTTGAATA      780
            *************** ********************************************

IS26.3      AGACAAAAGGCTGCCTCATCGCTAACTTTGCAACAGTGCC  820
IS26.1      AGACAAAAGGCTGCCTCATCGCTAACTTTGCAACAGTGCC  820
IS26        AGACAAAAGGCTGCCTCATCGCTAACTTTGCAACAGTGCC  820
IS26.2      AGACAAAAGGCTGCCTCATCGCTAACTTTGCAACAGTGCC  820
            ****************************************
```

### Comparison between IS*5075* and IS*5075.1*

The sequence of IS*5075*.1 (query) identified in IncHI1 plasmids pEB3_IMP1, pIMP47, and pIMP76_1 was aligned to the sequence of IS*5075* (subject) in the ISfinder database using megaBLAST on the National Center for Biotechnology Information (NCBI) website (blast.ncbi.nlm.nih.gov/Blast.cgi). Dots represent identical bases in this alignment.

```
Program: BLASTN
Query: IS5075.1 ID: lcl|Query_2929567(dna) Length: 1327
Subject:IS5075 ID: lcl|Query_2929569(dna) Length: 1327

>IS5075
Sequence ID: Query_2929569 Length: 1327
Range 1: 1 to 1327

Score:2429 bits(1315), Expect:0.0,
Identities:1323/1327(99%),  Gaps:0/1327(0%), Strand: Plus/Plus

Query  1     TAATGAGATGGTCACTCCCTCCTTCCCAGTACTATGCTGAGGACAGGCTTTCATTCGGAG  60
Sbjct  1     ............................................................  60

Query  61    AACCATCATGGAAAACATTGCGCTTATTGGTATCGATCTGGGTAAGAACTCTTTCCATAT  120
Sbjct  61    ............................................................  120

Query  121   TCATTGTCAGGATCATCGTGGGAAGGCCGTTTACCGTAAAAAATTCACCCGACCAAAGCT  180
Sbjct  121   ............................................................  180

Query  181   AATCGAATTTCTGGCGACATGCCCGGCAACAACCATCGCGATGGAAGCCTGTGGCGGTTC  240
Sbjct  181   ............................................................  240

Query  241   TCACTTTATGGCACGCAAGCTGGCAGAGTTAGGGCATTTTCCAAAGCTGATATCACCGCA  300
Sbjct  241   ............................................................  300

Query  301   ATTTGTCCGCCCATTCGTTAAAAGCAACAAAAATGACTTCGTTGATGCTGAAGCTATCTG  360
Sbjct  301   ............................................................  360

Query  361   TGAAGCAGCATCACGTCCATCTATGCGTTTCGTGCAGCCCAGAACCGAATCTCAGCAGGC  420
Sbjct  361   ............................................................  420

Query  421   AATGCGAGCTCTGCATCGTGTCCGTGAATCCCTGGTTCAGGATAAGGTGAAAACAACTAA  480
Sbjct  421   ............................................................  480

Query  481   TCAGATGCATGCTTTTCTGCTGGAATTTGGTATCAGCGTTCCGCGAGGTGCTGCCGTTAT  540
Sbjct  481   ............................................................  540

Query  541   TAGTCGACTGAGTACCCTTCTTGAGGACAGTAGTTTGCCTCTTTATCTCAGCCAGTTACT  600
Sbjct  541   ................G..............C............................  600

Query  601   GCTGAAATTACAACAGCATTATCACTATCTTGTTGAGCAGATTAAAGATCTGGAATCTCA  660
Sbjct  601   ............................................................  660

Query  661   GTTGAAACGAAAGTTGGACGAAGATGAGGTTGGACAGCGCTTGCTGAGTATTCCCTGCGT  720
Sbjct  661   ............................................................  720
```

```
Query  721   TGGAACGCTGACTGCCAGTACTATTTCAACTGAGATTGGCGACGGGAAGCAGTACGCCAG  780
Sbjct  721   ............................................................  780

Query  781   CAGCCGTGACTTTGCGGCGGCAACAGGGCTGGTACCCCGACAGTACAGCACGGGAGGTCG  840
Sbjct  781   ............................................................  840

Query  841   GACGACATTGTTAGGGATTAGCAAGCGGGGCAACAAAAAGATCCGAACTTTGTTGGTTCA  900
Sbjct  841   ............................................................  900

Query  901   GTGTGCCAGGGTATTCATACAAAAACTGGAACACCAGTCTGGCAAGTTGGCCGACTGGGT  960
Sbjct  901   ............................................................  960

Query  961   CAGGGAGTTGTTGTGTCGGAAAAGCAACTTTGTCGTCACCTGTGCTCTGGCAAACAAGCT  1020
Sbjct  961   ............................................................  1020

Query  1021  GGCCAGAATAGCCTGGGCACTGACGGCGCGACAGCAAACTTACGAAGCATAAAGGCAGAA  1080
Sbjct  1021  ............................................................  1080

Query  1081  ATACACCAGTTTAAACAATCATTCATCTGGTTTTGCGAATACTGATATTGATGATACTAA  1140
Sbjct  1081  ............................................................  1140

Query  1141  CGGCCCACCGGCCTGTTGAGGAACCTGTAAAACGGAAAGGCTCATTGAAGCCGTATATTT  1200
Sbjct  1141  ............................................................  1200

Query  1201  TCTGGAGGTTCATCAGGCGCGGAACTCATCGAGGCGCGGGAATAAAATCCCATTCAGACG  1260
Sbjct  1201  ............................................................  1260

Query  1261  CCGGATAGATTCAAGCAAGCCAACTTGTCGTCAAAATCGGTGTTGCAAAAACGGGAGTGA  1320
Sbjct  1261  .......................T..A.................................  1320

Query  1321  CCATAGA  1327
Sbjct  1321  .......  1327
```
